# *In-vivo* Therapeutic Evaluation of the Wound Healing: Potential of Ethanolic Extract of *Trichilia heudelotii* Leaves in Wistar Rats

**DOI:** 10.1101/2023.12.14.571768

**Authors:** Awolola Gbonjubola Victoria, Jinadu Taiwo Hassan, Yakubu Musa Toyin, Oyewopo Adeoye Oyetunji, Abdullahi Tunde Aborode, Onifade Isreal Ayobami

## Abstract

**Background:** *Trichilia heudelotii* belongs to the *Meliaceae* family and is commonly used in indigenous African medicine to prevent and treat various microbial ailments.

**Methods:** *Trichilia heudelotii’s* ethanolic leaf extracts contains phytochemicals which was studied to demonstrate its wound-healing properties. Phytochemical screening of leaf extracts contains flavonoids, phenolics, tannins, terpenoids, saponins, and alkaloids, which play a vital part in the healing process. Overall, quantitative, and qualitative screening were performed to evaluate the therapeutic efficacy of *Trichilia heudelotii’s* phytochemicals for wound treatment.

**Results:** The wound-healing potency was evaluated using different concentrations of formulated oil extracts (0.5, 1.0, and 2.0g/10g) on fresh wounds inflicted on Wistar rats using the excision model. Healing capacity was assessed based on rate of wound contracting ability, time of wound closure, and epithelisation period. The results shows that leaf extract oils exhibited significant wound healing capacity (p<0.05) compared with controls. On the 18th day in the infected group using 2.0/10g extract formulated ointment, 100% wound closure was observed. On day 21, using 0.5/10g and 1.0/ 10g, extract formulated ointment compared with controls which exhibited 79% and 89%, respectively. In addition, histopathological characteristics show increased and well-organized new bands of collagen and fibroblasts in the extract formulated treatment than in the controls, which supports the wounding healing potential of the plant. This is the first report providing therapeutic evidence for the wound healing activity of ethanolic leaf extract on Wistar rats.

**Conclusion:** This indicate the evidence that establish the traditional use of *Trichilia heudelotii* extract for treating skin infections and wounds. The results also raise the possibility of developing a commercialized medicinal product-based venture based on this extract.

**Graphical Abstract:** 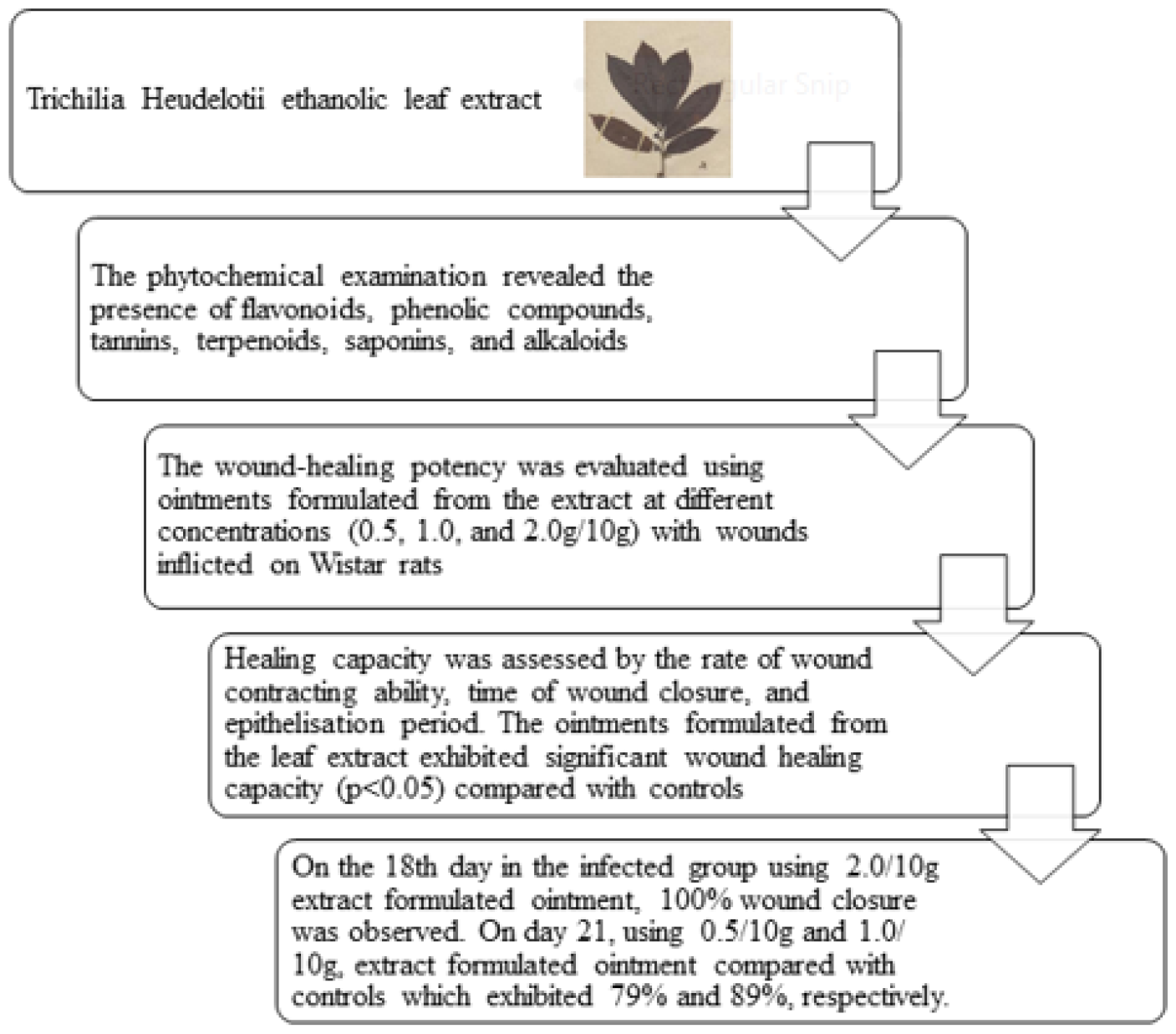

## Background

Injuries that tear or damage the skin are considered wounds because they alter the structure and composition of the epidermis (Strodtbeck, 2001). Wound healing is the process through which injured skin cells is repaired (Sherratt and Dallon, 2002). Antibiotics, irrigation, cleaning, tissue transplants, and proteolytic enzymes are being employed in the several wound treatment and management practices (Nayak et al., 2007). Depending on the variety, the medium-sized tree *Trichilia heudelotii* Planch (Meliaceae) can reach heights of 12-20 meters and is referred to in Yoruba as “Akorere,” “Akokorere,” “Aka,” or “Akika” (Solomon et al., 2005). Terms of location-wise, it is available in the southwestern part of Nigeria and the tropical rainforest region of Africa. Diverse bacterial illnesses, including those of the digestive tract and skin and wounds, are treated with the plant through its ethnobotanical uses (Nilugal et al., 2017; Nwala et al., 2013; Adeniyi et al., 2008; Nayak et al., 2007). *Trichilia heudelotii* bark may be effective for treating various medical conditions, including but not limited to coughing, syphilis, and skin disease (Abbiw, 1990; Irvine, 1961). Historically, the stem bark has served medicinal purposes, including as a broad-spectrum antimicrobial, stimulant, emergency contraceptive, and anti-plasmodial (Abbiw, 1990). *Trichilia heudelotii* ethanolic extracts have been shown to have antimicrobial activities, and this activity has been linked to dose content (Bankole et al., 2015). The antibacterial activity of methanol extracts of *Trichilia heudelotii* leaf has been documented (Aladesanmi and Odediran, 2000). A phytochemical called *heudebolin* has been isolated from the root extract (Okorie and Taylor, 1967). The n-hexane extract of the leaves of *Trichilia Heudelotii* yields diterpenes, which include nimbiol, 7-ketoferruginol, isopimarinol, and 12β-hydroxysandaracopimar-15-ene, while the ethyl acetate extract yield protocatechuic acid, 4-hydroxybenzoic acid, 2-methylproto-catechuic acid and 2-propionoxy-β-resorcylic acid (Aladesanmi and Odediran 2000). The plant is a source of food in cases of low blood sugar in children and pregnant women, as well as in cases of injury and wounds (Aladesanmi and Odediran, 2000). Another unproven folklore claim is that *Trichilia heudelotii* leaves can be used to cure wounds. However, there has been little to no research on the wound-healing ability of *Trichilia Heudelotii; thus*, this study aims to assess the phytoconstituents makeup of the plant and its promise for use in the treatment of wounds.

## Methods

### *Trichilia heudelotii* leaves

#### Sample collection and Identification

The leaves of *Trichilia heudelotii* were obtained at Apaara village at Oyo State, Nigeria, and was established for documentation in the Herbarium Unit of the Department of Plant Biology, University of Ilorin, Nigeria, where the voucher number of UILH/001/1367 was given to the plants identified.

#### Sample preparation and extraction

The leaves of *Trichilia heudelotii* were sun dried under the tree, however, before the chemical analysis begins, around 500grams of the sun dried leaves were crushed with mechanic grinder and the extracts from the grinder were mixed with ethanol for around 48 hours. However, to obtain the crude sample after the ethanol mixture, filtration and evaporation takes place.

### Qualitative screening of leaves phytochemicals

#### Determination for Tannins

A small sample of 1% of lead acetate was mixed with 0.2gram of the extract obtained and the reaction was noticed for a while till the sample form a precipitate with a yellow colour (Trease and Evans, 1989).

#### Determination for Alkaloids

The 0.2g of the leaf extracts was added to 5ml of aqueous 1% of HCl, the solution was put inside the water bath for a while before it undergo filtration. The obtained results from the filtration was categorized into two test tubes by placing 1ml samples of the filtrate obtained into each of the test tubes. After the division of the filtrate into two part, Mayer’s reagent is added to the first part of the test tube which contain the first filtrate, a precipitation is obtained with a buff colour that indicate the presence of the alkaloids. While in the second part of the test tubes, a small portion of solution of Dragendorff’s was mixed with the filtrate, a precipitation is obtained with a orang-red colour noticed (Sofowora, 1993).

#### Determination for Terpenoids (Salkowski test)

Around 0.2g of the leaf extract was added to 2ml of chloroform, and 3ml of concentrated H_2_SO_4_ was mixed with the extract, after the mixture, there is an observation of reddish-brown colour at the surface of the sample that indicate the presence of terpenoids (Sofowora, 1993).

#### Determination for Phenolic Compounds

Sofowora (1993) revealed adding 0.5g of the leaf extracts to distilled water of 5ml, and a small portion of neutral ferric chloride solution of 5% will lead to dark-green colour, which identify the phenolic compounds presence in the extract.

#### Determination for Flavonoids

The part of the leaf extract was added to dilute Ammonia solution of 4ml in measurement and concentrated sulphuric acid, an observation of flavonoids is noticied due to the show of the yellow colour (Sofowora, 1993).

#### Determination for Cardiac Glycosides (Keller-Killiani test)

Ayoola et al. (2008) suggested that mixing 0.2g of the leaf extracts with glacial acetic acid of 2ml in measurement and one drop of ferric chloride solution, with a mixture with 1ml of concentrated H_2_SO_4_, this mixture result to observation that there is presence of cardiac glycosides with a show of brown ring on the surface of the mixed sample.

#### Determination for Saponins

Around 1g of the leaf extract was heated with distilled water of 5ml and after heating, the mixture under filtration, and the filtrate obtained was added with distilled water of about 3ml in measurement and was shaken for around five minutes till an observation of frothing was obtained that confirmed the saponins presence (Trease and Evans, 1989).

#### Determination for Anthraquinones

Around 0.2g of the leaf extract was mixed with a small portion of dilute sulphuric acid, and the mixture was heated, after heating, it under filtration. Also, the filtrate that was obtained from the filtration was mixed with the chloroform of 2ml in measurement and the surface of chloroform was mixed with 1ml of ammonia solution, the mixture identified the presence of anthraquinones with the formation precipitate and red colour (Sofowora, 1993; Harborne, 1973).

#### Determination for Phlobatannins

0.3g of the leaf extracts was heated with 1% of the aqueous hydrochloric acid, this mixture result to the a precipitation with red colour indication that established the phlobatannins (Sofowora, 1993; Harborne, 1973).

#### Determination for Coumarins

Savithramma et al. (2011) suggested that when a small portion of the leaf extracts undergo mixture with 10% of NaOH with 3ml in measurement, an observation of Coumarins presence was established through yellow colour formation (Savithramma *et al*., 2011).

#### Determination for Steroids

Sofowora (1993) suggested that when a small part of the leaf extracts mixed with 2ml Acetic anhydride and 2ml Sulphuric acid, it results to the formation of transitive colour change from violet to blue or green that indicates steroids presence.

#### Determination for Anthocyanins

In order to treat a portion of the leaf extract, 3ml of a 2M hydrochloric acid and ammonia solution were combined. Anthocyanins are present if pink-red solution forms, which thereafter turns blue-violet (Savithramma *et al*., 2011).

### Quantitative Determination of Phytochemicals

#### Determination of Tannins Content

In a bottom flask, 5g of leaf extracts were measured and mixed with 100ml of 2M HCl. The mixture was simmered for 30 minutes in a water bath, cooled, and then filtered. The filtrate was then obtained twice with 40ml of diethyl ether, and the ether isolate was dried and measured (Okwu and Iroabuchi, 2004).

#### Determination of Alkaloids content

A 250 ml beaker containing 5g of the leaf extracts was measured, the leaf extracts measured was mixed with 200ml of 10% acetic acid in ethanol, and left to stand for 4 hours with the lid tightly closed. The solution was then filtered, and the filtrate was then condensed in a water bath to 75 percent of its initial volume. The alkaloids were then precipitated by adding concentrated ammonia solution to the saturated extract dropwise. The resultant solution was given time to settle, then the precipitate was taken out and measured (Harborne, 1973).

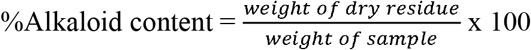

#### Determination of Total Terpenoids content

In 50ml of 95% aqueous ethanol, 2g of the material was bubbled up for 24 hours. After filtering the resultant mixture, petroleum ether was used to remove the filtrate, which was then dried out by concentration. Absolute terpenoids were determined by weighing the dried ether extracts (Harborne, 1973).

#### Determination of Total Phenolics content

N-hexane was used to dissolve 2g of leaf extract for 4 hours. To remove the fat content, the mixture was filtered, and the process was performed again on the residual. Diethyl ether was used to extract the powdered gelatin residual (sample) after which distilled water and 10% NaOH solution were added to the extract in a buchner funnel. Next, a 10% HCl solution was added to the separated water phase to acidify it to a pH of 4. The final extraction of the sample included the use of 50ml of dichloromethane, and later was recovered, dried, and used to measured the organic layer (Oluwaniyi and Oladipo, 2017).

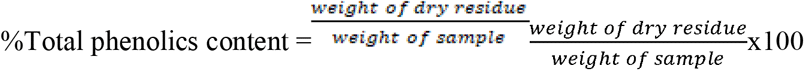

#### Determination of Flavonoids

At room temperature, 10g of the sample was periodically extracted with 100ml of 80% aqueous methanol. The resulting solution was then filtered. The filtrate was put into a beaker, dried over a boiling water, and weighed at a regular pace (Bohm and Kocipai-Abyazan, 1994).

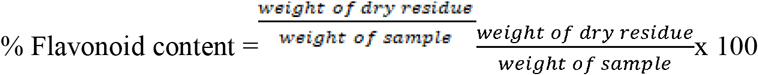

#### Determination of Saponins content

100ml of 20% ethanol were mixed with 5g of the sample. In a hot water bath at 55°C, the solution was heated and constantly agitated for 4 hours. Using Whatman No. 1 filter paper, the mixture was filtered, and the residue was extracted once more using an additional 100ml of 20% ethanol. A water bath heated to around 90°C was used to condense the combined extract to about 40ml. The concentration rinsed in diethyl ether before being extracted in n-butanol, which was then washed in 5% aqueous sodium chloride. The leftover solution was first boiled in a water bath before being dried to a consistent weight in the oven. A percentage was determined for the saponin content (Obadoni and Ochuko 2001).

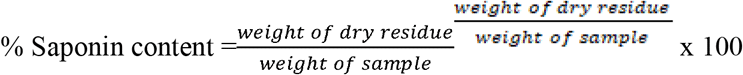

### Animals

For these in-vivo experiments, healthy male Wistar rats (weighing 180–200g) were employed. They were kept in regular laboratory settings in 24–280°C rooms with access to clean water and supplement-nourished meals (Amo Byng Nigeria Limited, Awe, Oyo State, Nigeria). The University of Ilorin Ethical Review Committee authorized all study designs used in this study, including those involving animals (Protocol Identification Code: UERC/PHY/154; UERC Approval Number: UERC/ASN/2018/1485) and conducted the research in accordance with Helsinki Declaration.

### Preparation of extract ointments

For this investigation, three sets of the extract ointments were made, each of which contained extract in various amounts (0.5g, 1.0g, and 2.0g per 10g of the ointment base). First, for each batch, 10g of petroleum jelly B.P. were put in a beaker and melted over a thermostatically controlled water bath. The needed amounts of the extracts were then measured out, added to the hot ointment base, and homogenized. Finally, the ointments were kept in the fridge until usage (Nwala *et al*., 2013).

### Infliction of wounds

The animals’ back hairs were removed by shaving. Under diethyl ether anesthesia, an excision wound with a total thickness of 314 mm^2^, a 20 mm diameter, and a 2 mm depth was made. The dorsal thoracic region of the anesthetized rat was imprinted 10 mm from the vertebral column and 50 mm from the ear. By dabbing the wound with a cotton swab soaked in regular saline, hemostasis was accomplished (Kiran et al., 2017).

In this study, five distinct groups of five animals each received the following topically administered, once day, treatment:

**Group I**- Animals that received injury for wound formation and treated with the blank ointment (Control).

**Group II**- Animals that received injury for wound formation and treated with the reference drug, i.e., Gentamycin cream (Standard).

**Group III**- Animals that received injury for wound formation and treated with 0.5g of extract/10g of ointment base (Test 1).

**Group IV**- Animals that received injury for wound formation and treated with 1.0g of extract/10g of ointment base (Test 2).

**Group V**- Animals that received injury for wound formation and treated with 2.0g of extract/10g of ointment base (Test 3).

### Measurement of wound area and wound contraction

The progressive changes in the wound area were monitored on days 0, 3, 6, 9, 12, 15, 18, and 21. In addition, wound areas were measured (Nwala *et al*., 2013). Wound contraction was calculated as a percentage of the reduction in the original wound area size (Preeti *et al*., 2017) using the formula:

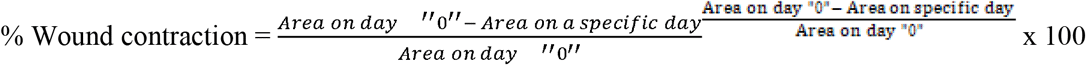

### Histopathological analysis

At the end of the experiment, a tissue sample was taken from the skin of each set of rats and analyzed for histopathological changes. After being processed and blocked with paraffin, the specimens were fixed in 10% buffered formalin, sectioned into 5mm, and stained with hematoxylin and eosin (H&E). These histological alterations, such as fibroblast growth and collagen production, were then examined under a microscope (Kiran *et al*., 2017).

### Statistical analysis

Version 16.0 of SPSS software was used to conduct the statistical analysis. All findings were presented in terms of mean and standard deviation. Comparative analyses between the treatment and control groups were done statistically. One-way ANOVA was used to analyze the wound area and wound contraction % data. A value of P < 0.05 was statistically significant (Steel *et al*. 1996).

## Results

### Qualitative and Quantitative Phytochemical Screening

**Table 1, the** Qualitative and Quantification phytochemical analysis of *Trichilia heudelotii* ethanolic leaf extract, is given below. Table one shows the quality and quantity analysis of *Trichilia heudelotii* ethanolic leaf extract. In the table, twelve phytochemicals were analyzed, and six were observed in the section. These include tannins, alkaloids, terpenoids, saponins, flavonoids, and phenolics. The six observed absent have anthraquinones, phlobatannins, coumarins, cardiac glycosides, steroids, and anthocyanins.

**Table 1:** Qualitative and Quantification phytochemical analysis of *T. heudelotii* ethanolic leaf extract.

| Phytochemicals |  | % Composition<br>(g/100g) |
| --- | --- | --- |
| Tannins | + | 6.31 ± 0.09 |
| Alkaloids | + | 0.98 ± 0.04 |
| Terpenoids | + | 3.45 ± 0.41 |
| Anthraquinones | - | - |
| Phlobatannins | - | - |
| Coumarins | - | - |
| Saponins | + | 2.86 ± 0.12 |
| Flavonoids | + | 13.77 ± 0.05 |
| Cardiac Glycosides | - | - |
| Steroids | - | - |
| Phenoilic Compounds | + | 9.02 ± 0.08 |
| Anthocyanins | - | - |

**Key:** +: present, -: absent

### Wound-Healing Activity

Wound healing time following resection in rats treated with ointments containing *Trichilia heudelotii* alcoholic leaf extract is shown in **Table 2**. Total wound closure occurred in the 2.0g/10g ointment large set on or before day 18 of the research (**Table 2; Figures 1**). Both groups receiving ointment base concentrations of 0.5g/10g and 1.0g/10g could achieve complete wound contraction by the end of the research (21 days). Total wound closure occurred in the gentamycin cream group on or before day 15 of the investigation. Although the control group’s wound size (32.97±11.73mm^2^) was negligible compared to the initial wound size (314 mm^2^), it was still present on day 28.

**Table 2:**
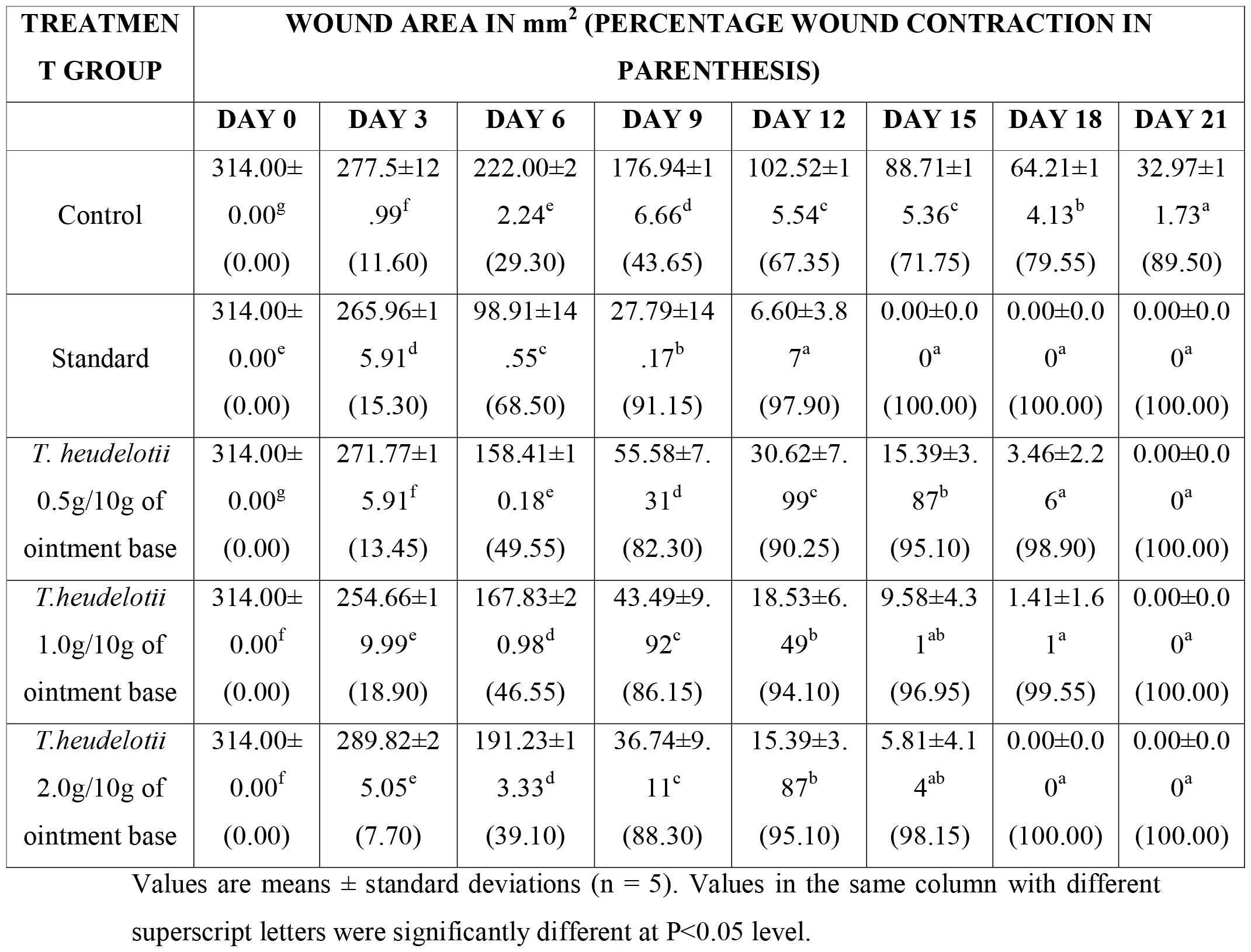
Effect of *Trichilia heudelotii* ethanolic leaf extract ointments on excision wound healing in rats.

**Figure 1:**
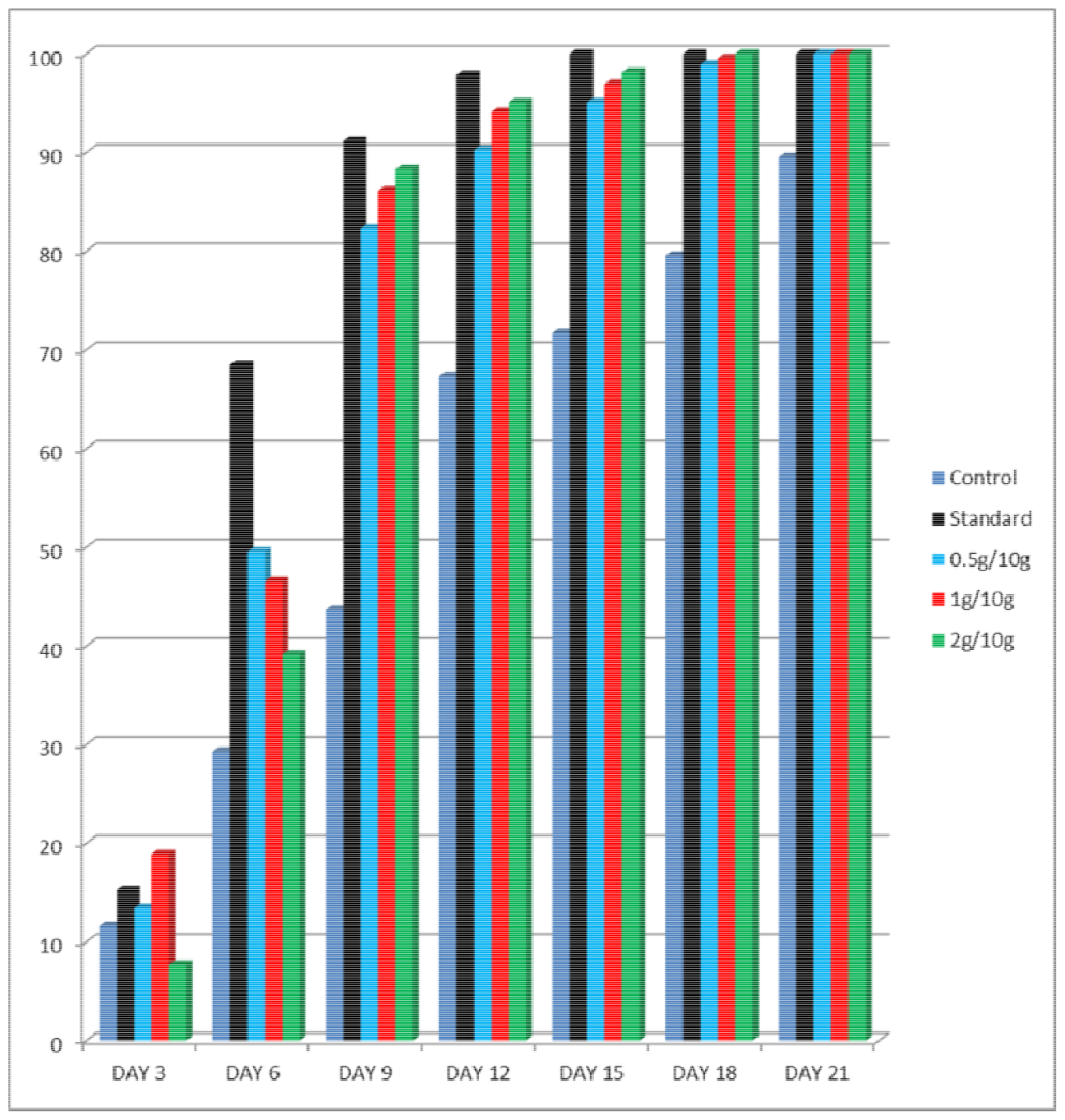
Percentage of wound contraction in relation to days (wound closure)

### Histopathological examinations

Figure 2. shows considerable wound healing as collagen fibers and fibroblast cells in albino rats treated with basic and topical formulations (0.5, 1.0, and 2.0g/10g of ointment base) (Plates 2, 3, 4, and 5, respectively). Cellular senescence of collagen fibres (incomplete healing) and fibroblast cells were observed in the wounds of the control group (Plate 1). There was a significant difference in the wound-healing activity between the groups treated with extract ointments and the group that used the control ointment alone. Collagen fibres and fibroblast are more visible in the regular and extracts cream treated groups, as compared to the control group.

## Discussion

The analysis revealed the amount of the flavonoids content to be the highest (13.77 ± 0.05g/100g) in *Trichilia heudelotii*, followed by total phenolics (9.02 ± 0.08g/100g), tannins (6.31 ± 0.09g/100g), total terpenoids (3.45 ± 0.41g/100g) and saponins (2.86 ± 0.12g/100g). The alkaloid content (0.98 ± 0.04g/100g) was the least.

**Figure 2:**
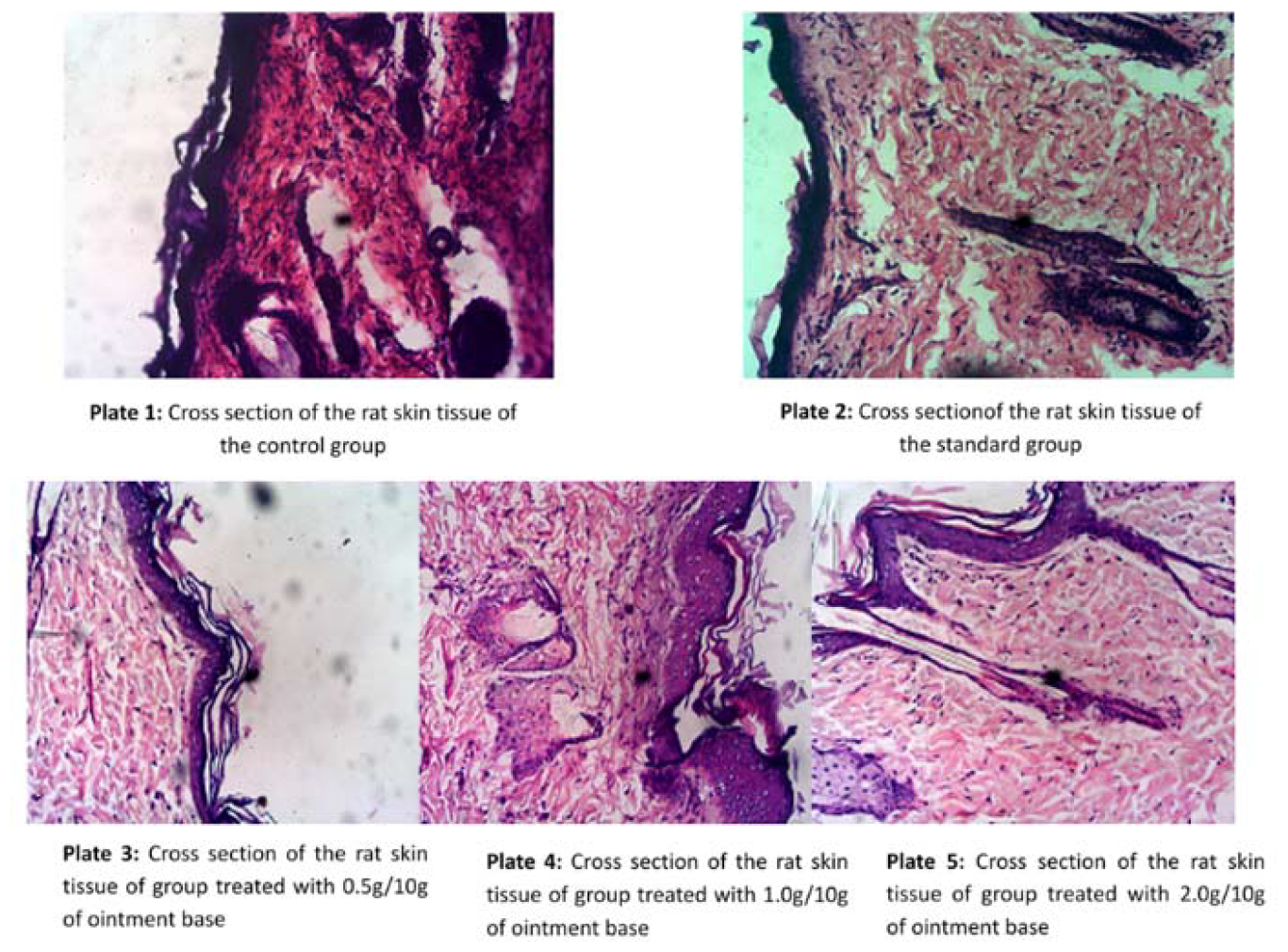
Histopathological examinations

*Trichilia Heudelotii* has a wide range of therapeutic effects, including antibacterial activity, and the ethanolic leaves extract contains the active compounds responsible for these effects, including alkaloids and saponins (Ngbolua, 2014; Idu, 2012). Because of their natural tendency to repel microorganisms, microbes are promising options for treating illnesses, as stated by Okwu (2006). Antimicrobial chemicals like saponins, alkaloids, and others like them play an essential role in the body’s defense against diseases and other microorganisms (Okwu, 2005).

Flavonoids were also found in the analysis, and their astringent and antibacterial qualities are estimated to be associated with wound closure and an accelerated rate of epithelialization, both of which have been linked to improved wound healing (Tsuchiya et al., 1996; Scortichini and Pia, 1991). The observation supports the antibacterial and antioxidant actions of plants that tannins have this effect (Kar, 2007).

Therefore, *Trichilia heudelotii’s* discovered compounds, such as triterpenoids, alkaloids, and flavonoids, may play an essential part in the recovery process. So, numerous research has done a variety of things to determine if *Trichilia heudelotii* is effective at healing wounds (Kar, 2007; Ngbolua, 2014; Idu, 2012).

The phytochemical screening assay is an example of this procedure. Tannins, alkaloids, lactate dehydrogenase, steroids, saponins, terpenoids, and flavonoids are all found in the methanol extracts, as shown by the phytochemical screening (Idu, 2012). Further phytoconstituents investigations are necessary to isolate the effective compound(s) willing to take responsibility for these pharmacological effects; however, it is possible that the components of the extract of leaves, such as terpenoids, flavonoids, and alkaloids, play an essential role in the wound healing observed in the present study (Nayak et al., 2009).

Astringent and antibacterial characteristics of terpenoids, flavonoids, and alkaloids are important for wound contraction and an enhanced rate of epithelialization two hallmarks of the wound healing process (Scortichini et al., 1991). Terpenoids, flavonoids, and alkaloids are well-known compounds that, depending on their actions on the non-mevalonate route, may have significant antifungal or antimicrobial activity.

Fungal, protozoan, gram-negative bacterial, and other microbes use this process to produce cell membrane components, prenylation proteins, and a secondary carbon source (Nayak et al., 2010). Similar phytochemical elements, essential for increasing wound healing activity in rats, have been found in the study with different plant materials (Nayak et al., 2006).

On/before day 18, the rats in the groups treated with the gentamycin cream and 2.0g/10g of ointment base had observed complete wound healing (100.00% wound contraction) (**Table 2**; **Figures 1 and 2**). On/before the last day of the study (21 days), the groups treated with 0.5g/10g of ointment base and 1.0g/10g of ointment base showed complete wound healing (100.00% wound contraction).

In Figure 2 above, the analysis revealed that the standard had a more rapid wound healing response followed by 2.0g/10g, 1.0g/10g, and 0.5g/10g ointments. The control was found to have the least healing response. Comparatively, in Table 2, the treated 314 mm^2^ excision wounds revealed that the extracts had a healing ability ranging from 10.50 – 20.45% more than the control on days 18 to 21. The test drug achieved a higher and better healing difference of 28.25%, 4.90%, 3.05%, and 1.85% compared to the control’s healing abilities, 0.50g, 1.00g, and 2.00g leaf extract concentrations on day 15 of the analysis.

However, the leaf extract healing ability of 2.00g in 10.00g base concentration was compared favourably with the standard test drug on day 15. They had an insignificant healing difference of 1.85%, which shows that the leaf extract concentration of 2.0g/10g had a better healing response than other lower concentrations whose healing difference as compared with the standard on day 15 were 4.90% and 3.05% for 0.50g/10g and 1.00g/10g concentrations respectively.

As a result, it is reasonable to assume that a higher concentration of extract of leaves in the foundation used for balm compositions will result in a more rapid recovery process.

One can deduce from the plates that the leaf extract helped the resection wound recover more quickly than the control group. Therefore, the extract ointments have a noticeable pro-healing effect, as evidenced by the large rise in the rate of expansion. Thus, the findings of this study provide scientific support for the use of Trichilia heudelotii leaf extracts in wound care and healing.

## Conclusions

The research study shows that the ethanolic leaf extracts of *Trichilia heudelotii*, formulated into a simple ointment base, have achieved significant wound-healing activity. The various secondary metabolites present in this leaf ethanolic extracts should be harnessed to treat various skin wounds. More research should be carried out to ascertain which of the phytochemicals is responsible for a more healing property from the leaf of this plant.

## Abbreviations

NaOH: Sodium hydroxide
H_2_SO_4_: Sulfuric acid
HCl: Hydrochloric acid
(H&E): Haematoxylin and Eosin

## Ethics approval and consent to participate

All research protocols in our study, including animals, were approved by the University of Ilorin Ethical Review Committee (Protocol Identification Code: UERC/PHY/154; UERC Approval Number: UERC/ASN/2018/1485) and conducted in accordance with Helsinki Declaration. There were no humans’ participants.

## Consent to participate

Not Applicable

## Consent to publish

Not Applicable

## Availability of data and materials

Not Applicable

## Acknowledgment

Taif University Researchers Supporting Project number (TURSP-2020/38), Taif University, Taif, Saudi Arabia.

## Authors’ contributions

AGV, JTH, YMT, OAO, ATA: Experimental work and manuscript writing; MP, MHR, GSB, AA, EAF: Conceptualization, study design, and analysis; SSA, SMA, NM, SM: Critical Revision on the final manuscript and Funding Acquisition. All authors approved the final version of the manuscript.

